# Host immune stress reveals a role for potassium homeostasis in *Pseudomonas aeruginosa* aggregate stability

**DOI:** 10.64898/2026.09.09.750359

**Authors:** Oriana M. Williams, Sophie E. Darch

## Abstract

*Pseudomonas aeruginosa* (*Pa*) forms multicellular aggregates during chronic airway infection, yet the physiological processes that regulate aggregate organization and dispersal remain incompletely understood. Here, we investigated how *Pa* aggregates respond to human neutrophil elastase (HNE), a host-derived protease abundant in the cystic fibrosis airway. Transcriptomic analysis of HNE-exposed aggregates identified strong induction of the kdpFABC high-affinity potassium transport system, with greater induction in aggregate than planktonic populations. Functional analysis of *kdpA* revealed a broader role in aggregate physiology: loss of kdpA increased total biomass while reducing average aggregate volume and accelerating dispersal, including in the absence of HNE. Genetic complementation restored key *kdpA*-dependent phenotypes, whereas co-culture with wild-type cells did not rescue the mutant. Increasing extracellular potassium modified aggregate volume, biomass, and dispersal in the *kdpA* mutant but did not uniformly restore these phenotypes to wild-type levels, demonstrating that kdpA-dependent aggregate behavior is potassium-responsive. Together, these findings identify KdpA as a link between potassium homeostasis and *Pa* aggregate behavior and suggest that host-derived stress increases engagement of a homeostatic system that contributes to the coordination of population growth, aggregate organization, and dispersal.

## INTRODUCTION

Chronic *Pseudomonas aeruginosa* (*Pa*) infections remain a major clinical challenge in people with cystic fibrosis (pwCF). Within the CF airway, *Pa* encounters a dynamic environment characterized by oxygen limitation, nutrient variability, and sustained host inflammatory pressure(1–3). Importantly, *Pa* in the CF airway predominantly exists as free-growing multicellular aggregates rather than as surface-attached biofilms(4). This growth state is associated with distinct physiological properties and increased tolerance to antimicrobial and host-derived stresses(5, 6). However, the bacterial pathways that regulate aggregate physiology and enable adaptation to the inflammatory airway environment remain incompletely understood.

A hallmark of chronic CF airway disease is persistent neutrophilic inflammation(7). Neutrophils deploy multiple antimicrobial mechanisms, including phagocytosis, production of reactive oxidants, release of antimicrobial peptides, and formation of neutrophil extracellular traps(8, 9). Human neutrophil elastase (HNE), a serine protease released during neutrophil degranulation and NET formation, accumulates at high levels in CF airways and contributes substantially to tissue damage and chronic inflammation(10–14). HNE can also directly influence *Pa* behavior(15), including bacterial aggregation, suggesting that this host-derived protease may shape bacterial physiology as well as contribute to host tissue pathology. How aggregate populations physiologically respond to HNE, however, remains poorly understood.

Potassium is a major intracellular cation in bacteria and contributes to fundamental processes including osmotic homeostasis, intracellular pH regulation, and membrane-associated physiology(16, 17). Bacteria maintain potassium homeostasis through multiple transport systems, including the high-affinity KdpFABC transporter, which is regulated by the KdpDE two-component system in response to conditions that increase the requirement for potassium uptake(18, 19). Beyond these fundamental cellular functions, potassium has also emerged as a potential regulator of multicellular bacterial behavior. Potassium flux can mediate electrical signaling within bacterial communities(20), while *Pa* has recently been shown to respond to host-derived potassium through the Kdp system to influence attachment, coalescence, and biofilm formation(21). Whether potassium homeostasis similarly contributes to the organization and dynamic behavior of free-growing *Pa* aggregates remains unknown.

Here, we investigated the response of *Pa* aggregates to HNE and identified strong induction of the kdpFABC high-affinity potassium transport system. Functional analysis of *kdpA* revealed a broader role in aggregate physiology, with loss of *kdpA* altering population growth, aggregate organization, and dispersal both in the presence and absence of HNE. Genetic complementation restored key *kdpA*-dependent phenotypes, while manipulation of extracellular potassium demonstrated that aggregate behavior was potassium-responsive. Together, these findings identify KdpA as a link between potassium homeostasis and *Pa* aggregate behavior and support a model in which host-derived stress increases engagement of a homeostatic system that contributes to the coordination of population growth, multicellular organization, and dispersal.

## RESULTS

### Neutrophil elastase induces expression of the potassium transport system *kdpFABC*

To identify bacterial pathways associated with the aggregate response to host-derived stress, we performed RNA sequencing of *Pa* populations grown in synthetic cystic fibrosis sputum medium (SCFM2) and exposed to physiologically relevant concentrations of human neutrophil elastase (1X HNE [10 μg/L] and 2X HNE [20 μg/L]) (22). HNE exposure resulted in broad transcriptional changes, with an overall shift toward gene upregulation (Figure 1A). Among genes exhibiting ≥ +/-2-fold induction, a substantial proportion encoded hypothetical proteins. Functional classification of annotated genes revealed a broader representation of pathways at the higher HNE concentration, including pathways associated with stress responses and cellular homeostasis (Figure 1B).

**Figure 1.**
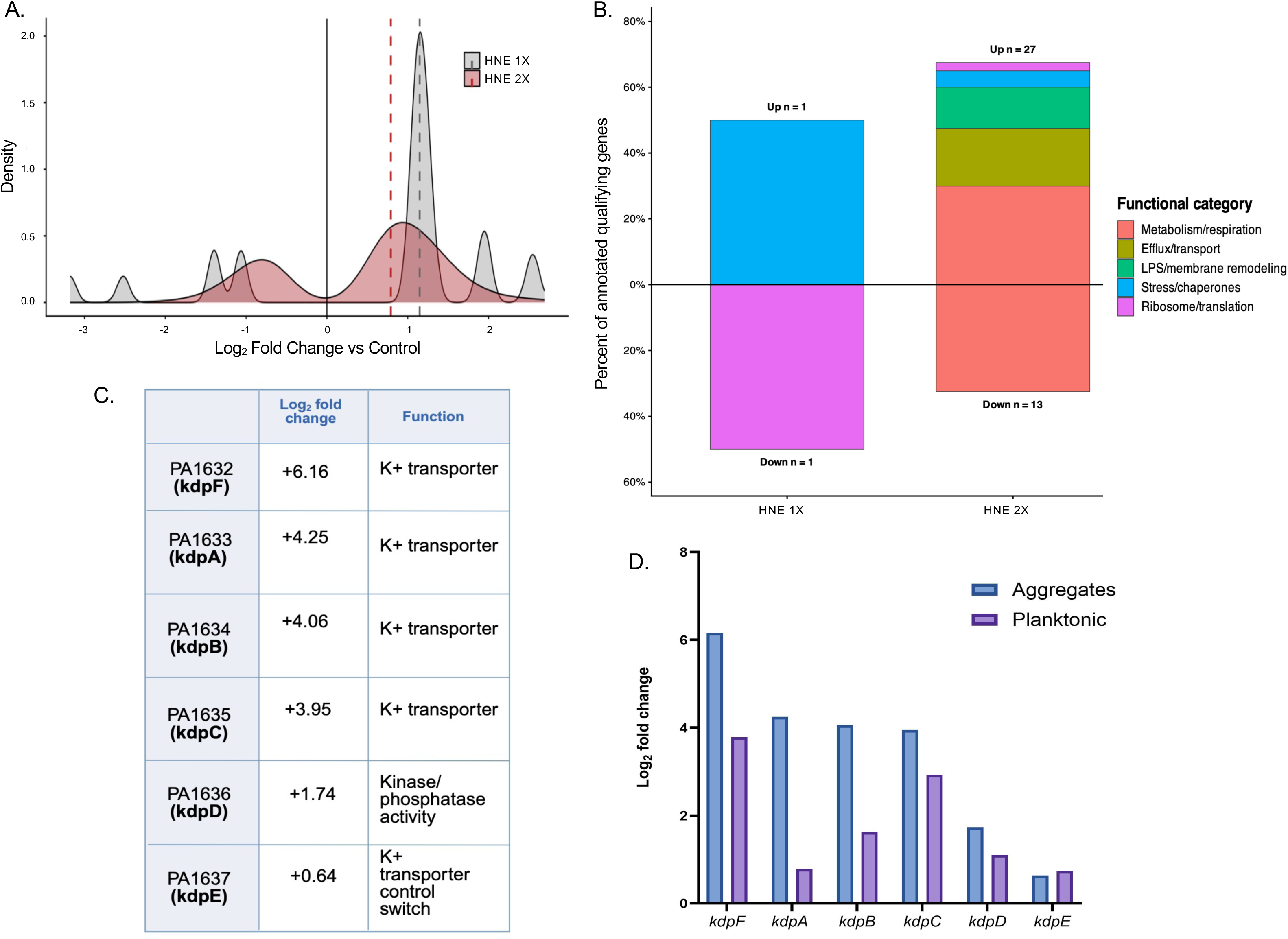
Human neutrophil elastase induces the Kdp potassium transport system in *Pa* aggregates. (A) Distribution of gene-expression changes in free-floating *P. aeruginosa* aggregates following exposure to 1X HNE (10 μg/L) or 2X HNE (20 μg/L), expressed as log_2_ fold change relative to untreated aggregates. (B) Distribution of functional categories among significantly differentially expressed genes (± log_2_ fold change) with annotated functions following exposure to 1X or 2X HNE. (C) Organization and function of components of the *kdp* operon, with corresponding log_2_ fold-change values in aggregates exposed to 2X HNE. (D) Comparison of log_2_ fold-change values for components of the *kdp* operon in aggregate and planktonic populations following exposure to 2X HNE.

Within this transcriptional response, we identified induction of the *kdpFABC* potassium transport system (Figure 1C). *kdpA*, encoding a component of the high-affinity Kdp potassium transporter, exhibited >4-fold induction in aggregates exposed to 2× HNE relative to untreated populations. Expression of genes within the *kdp* operon generally increased with increasing HNE exposure, although not all components met the predefined fold-change threshold for prioritization. Specifically, *kdpA*B (PA1633/PA1634) met the prioritization criteria in aggregate populations, whereas *kdpFC* (PA1632/PA1635) exhibited ≥2-fold changes in both aggregate and planktonic populations. The regulatory genes *kdpDE* (PA1636/PA1637) showed more modest increases and did not meet the prioritization threshold.

Although induction of the *kdp* operon was not exclusive to aggregate populations, the magnitude of the transcriptional response was greater in aggregates than in planktonic populations following HNE exposure (Figure 1D). Together, these findings identify potassium homeostasis as a prominent component of the aggregate transcriptional response to HNE and led us to investigate the functional contribution of KdpA to aggregate physiology.

### Loss of *kdpA* alters aggregate growth and organization

To determine whether KdpA contributes to aggregate physiology, we characterized a *kdpA* transposon mutant (PA1633::Tn) during growth in SCFM2. Loss of *kdpA* altered population growth dynamics, with the mutant exhibiting increased average biomass relative to wild type under untreated and HNE-exposed conditions (Figure 2A). Despite this increase in average biomass, the *kdpA* mutant formed smaller aggregates than wild type, with reduced average aggregate volume over the course of growth (Figure 2B). This phenotype was observed both in the presence and absence of HNE, indicating that the contribution of KdpA to aggregate organization is not restricted to HNE exposure.

**Figure 2.**
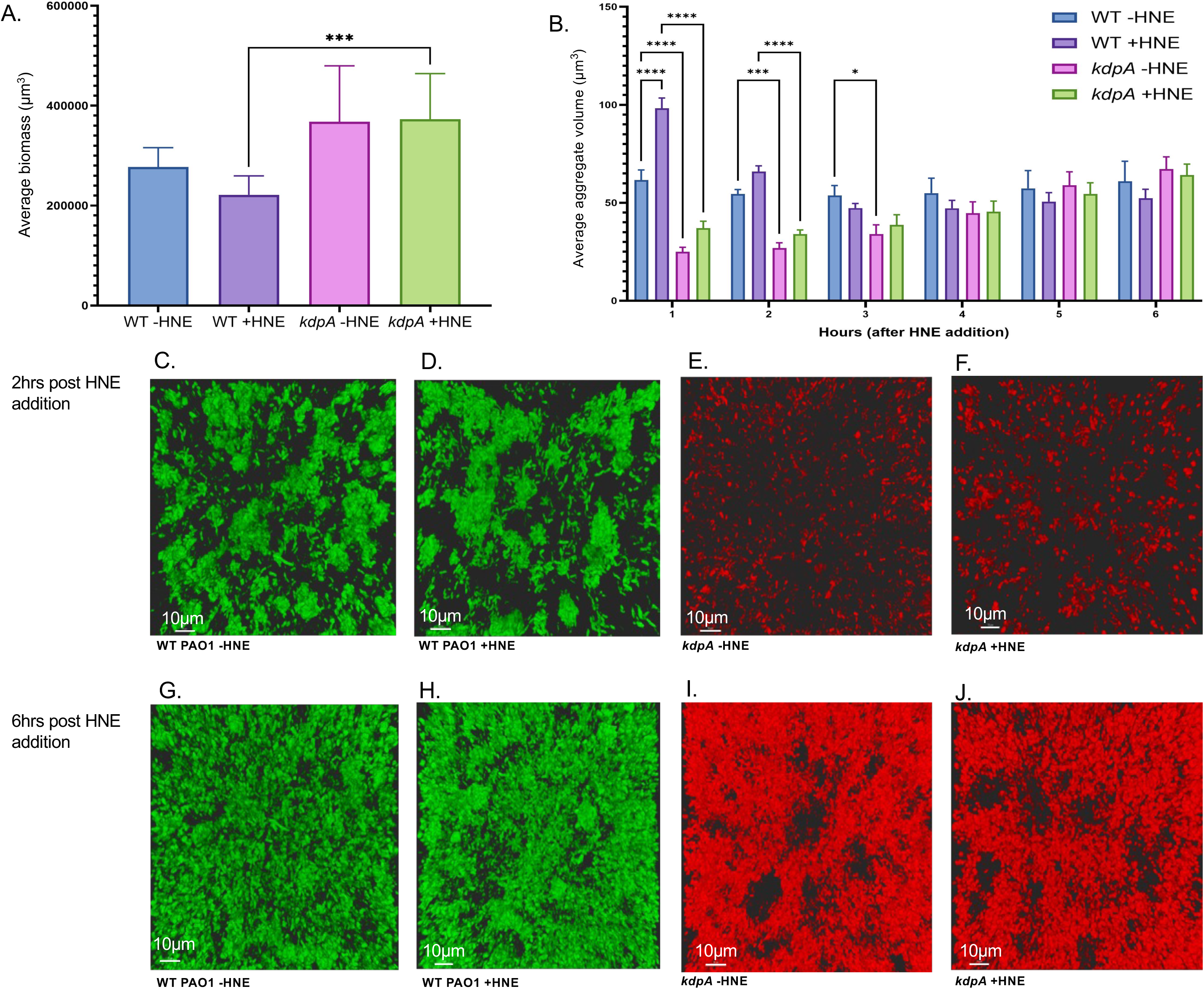
Loss of *kdpA* alters aggregate growth and organization. (A) Average biomass of WT and *kdpA* mutant aggregates grown in SCFM2 in the presence or absence of 2X HNE (20 μg/L). Data represent three independent biological replicates. Statistical significance was determined by two-way ANOVA with main effects followed by Šidák’s multiple-comparison test. Error bars are representative of ± SEM. (B) Average aggregate volume over time following HNE addition. Data represent three independent biological replicates. Statistical significance was determined by two-way ANOVA followed by Tukey’s multiple-comparison test. Error bars represent ± SEM. Statistical significance is indicated by an asterisk. (C–J) Representative CLSM images of WT GFP (C, D, G, H) and *kdpA* mutant mCherry (E, F, I, J) in the absence or presence of 2X HNE at 2 h (C–F) and 6 h (G–J) following HNE addition, corresponding to 6 h and 10 h total growth, respectively. Scale bars, 10 μm.

Confocal imaging further demonstrated differences in aggregate organization between wild-type and *kdpA* mutant populations (Figure 2C–J). At 2 h following HNE addition, mutant populations contained smaller aggregates than time-matched wild-type populations (Figure 2C–F). By 6 h, aggregates were apparent in both populations, but their organization remained visibly distinct (Figure 2G–J).

Together, these findings demonstrate that loss of *kdpA* alters the relationship between population growth and multicellular organization, resulting in increased biomass but reduced average aggregate volume. Thus, KdpA contributes to aggregate organization under basal aggregate-forming conditions as well as during HNE exposure.

### Loss of *kdpA* accelerates aggregate dispersal and alters membrane-associated DiBAC signal

Given the altered growth and architecture of *kdpA* mutant aggregates, we next examined whether loss of KdpA affected aggregate stability. Aggregate dispersal was quantified by measuring the number of planktonic (non-aggregating cells) over time in wild-type and *kdpA* mutant populations grown in SCFM2 in the presence or absence of HNE. The *kdpA* mutant exhibited earlier dispersal than wild-type populations (Figure 3A), indicating altered aggregate dispersal dynamics. At later time points, differences in dispersed cell numbers between mutant and wild-type populations became less pronounced as wild-type populations entered the dispersal phase. Within the *kdpA* mutant, HNE exposure did not significantly alter the number of dispersed cells over time, indicating that accelerated dispersal was primarily associated with loss of KdpA rather than being specifically induced by HNE.

**Figure 3.**
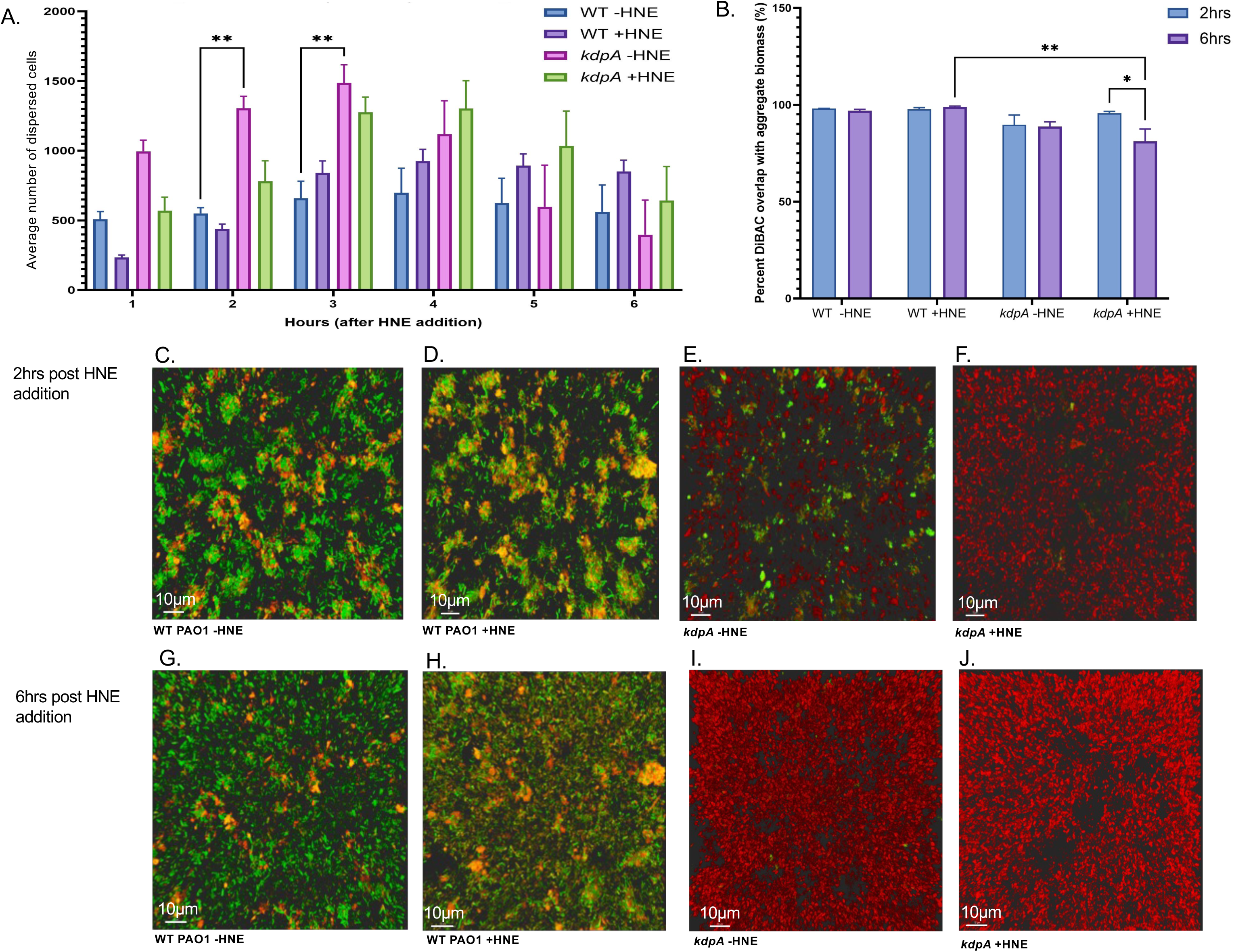
Loss of *kdpA* accelerates aggregate dispersal and alters membrane-associated DiBAC signal. (A) Average number of dispersed cells over time for WT and *kdpA* mutant populations grown in SCFM2 in the presence or absence of 2X HNE (20 μg/L). Data represent three biological replicates. Statistical significance was determined by two-way ANOVA followed by Tukey’s multiple-comparison test. Error bars represent ± SEM. (B) Percent overlap between detectable DiBAC signal and aggregate biomass at 2 and 6 h following HNE addition, corresponding to 6 and 10 h total growth, respectively. Colocalization was quantified in Imaris by determining the proportion of detectable DiBAC signal spatially associated with *P. aeruginosa* biomass. This measurement does not represent the percentage of DiBAC-positive cells. Data represent three independent biological replicates. Statistical significance was determined by two-way ANOVA followed by Šidák’s multiple-comparison test. Error bars represent ± SEM. Statistical significance is indicated by an asterisk. (C–J) Representative CLSM images of DiBAC signal associated with WT and *kdpA* mutant aggregates in the absence or presence of 2X HNE at 2 and 6 h following HNE addition. WT GFP was imaged with DiBAC4(5), and the *kdpA* mutant expressing mCherry was imaged with DiBAC4(3). Scale bars, 10 μm.

Because KdpA functions as a component of the high-affinity potassium transport system, we next asked whether loss of *kdpA* was associated with altered membrane-associated DiBAC signal. Wild-type and *kdpA* mutant aggregates were stained with spectrally compatible DiBAC dyes and imaged at 2 and 6 h following HNE addition (Figure 3C–J). WT GFP populations were imaged with DiBAC_4_(5), whereas the mCherry-expressing *kdpA* mutant was imaged with DiBAC_4_(3). Differences in detectable DiBAC signal were apparent between wild-type and mutant populations, with reduced detectable dye signal associated with *kdpA* mutant aggregates.

To quantify these differences, we measured colocalization between detectable DiBAC signal and aggregate biomass (Figure 3B). It is important to note that this analysis does not quantify the number or proportion of DiBAC-positive cells. Rather, colocalization represents the proportion of detectable DiBAC signal spatially associated with *Pa* biomass. Thus, differences between strains reflect changes in the association of detectable membrane-potential-sensitive dye signal with aggregate biomass rather than differences in the number of cells exhibiting DiBAC fluorescence.

Together, these findings demonstrate that loss of *kdpA* accelerates aggregate dispersal and is associated with altered membrane-associated DiBAC signal. Combined with the altered aggregate organization observed in Figure 2, these results support a role for KdpA in maintaining aggregate stability and are consistent with altered membrane-associated physiology during aggregate growth and HNE exposure.

### Genetic complementation restores *kdpA*-dependent aggregate phenotypes

To confirm that the altered aggregate phenotypes observed in the *kdpA* mutant resulted from disruption of *kdpA*, we generated a plasmid-based complementation strain expressing *kdpA*. Aggregate phenotypes were assessed at 6 h following HNE addition. Reintroduction of *kdpA* restored aggregate characteristics toward those observed in wild-type populations. At this time point, no significant differences were detected between the complemented strain and wild type in aggregate volume or dispersed cell numbers under either HNE condition (Figure 4A, C). These findings demonstrate that restoration of *kdpA* expression rescues key defects in aggregate architecture and dispersal associated with loss of KdpA.

**Figure 4.**
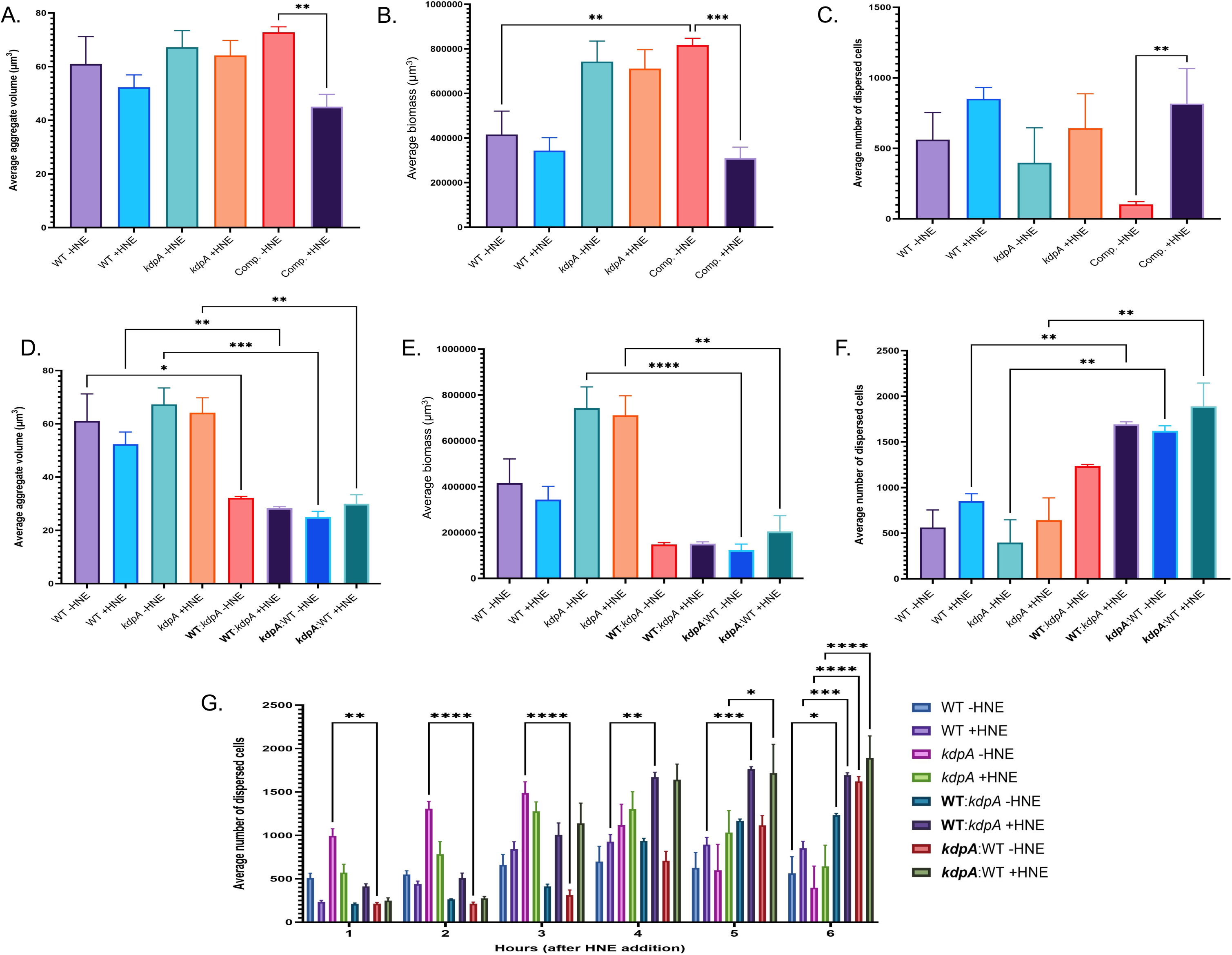
Genetic complementation restores *kdpA*-dependent aggregate phenotypes, whereas co-culture with wild-type cells does not rescue the mutant. (A–C) Average aggregate volume (A), average biomass (B), and average number of dispersed cells (C) for WT, *kdpA* mutant, and genetically complemented *kdpA* mutant populations (indicated by Comp.) at 6 h following addition of 2X HNE (20 μg/L). Data represent three biological replicates. (D–F) Average aggregate volume (D), average biomass (E), and average number of dispersed cells (F) for WT and *kdpA* mutant populations grown in monoculture or co-culture at 6 h following HNE addition. Analysis of individual strain when in co-culture is indicated in bold. Data represent two biological replicates. Statistical significance for A–F was determined using a Kruskal–Wallis test. Error bars represent ± SEM. (G) Average number of dispersed cells over time for WT and *kdpA* mutant populations grown in monoculture or co-culture following HNE addition. Data represent three independent biological replicates. Statistical significance was determined by two-way ANOVA followed by Tukey’s multiple-comparison test. Error bars represent ± SEM. Statistical significance is indicated by an asterisk.

We next asked whether the *kdpA* mutant phenotype could be rescued in trans by growth in the presence of wild-type cells. Wild-type and *kdpA* mutant populations were grown together and analyzed at 6 h following HNE addition. In contrast to genetic complementation, co-culture with wild-type cells did not restore mutant aggregate characteristics. Instead, both populations formed significantly smaller aggregates in co-culture than under their respective monoculture conditions, regardless of HNE exposure (Figure 4D). Biomass was also reduced during co-culture, with a significant reduction observed in *kdpA* mutant populations under both HNE treatment conditions (Figure 4E). These changes were accompanied by increased numbers of dispersed cells in co-culture compared with monoculture populations (Figure 4F).

To examine the spatial association of wild-type and mutant populations during co-culture, we quantified colocalization between the two populations at 2 and 6 h following HNE addition (Supplemental Figure 4). The two populations showed measurable spatial association within co-cultured aggregates, confirming that the absence of phenotypic rescue occurred despite physical association between wild-type and mutant cells.

Together, these findings demonstrate that genetic restoration of *kdpA* rescues key mutant aggregate phenotypes, whereas the presence of wild-type cells in co-culture does not. This lack of rescue in trans is consistent with KdpA functioning through a cell-associated process rather than through a diffusible factor supplied by neighboring wild-type cells.

### Potassium supplementation modifies *kdpA*-dependent aggregate phenotypes

We examined whether increasing extracellular potassium could compensate for loss of KdpA. WT and *kdpA* mutant populations were grown in SCFM2 supplemented with 25 mM KCl, and aggregate phenotypes were assessed in the absence and presence of 2X HNE (20 μg/L). Under non-stressed conditions, KCl supplementation altered aggregate phenotypes in both strains (Figure 5A–C). In the *kdpA* mutant, KCl increased average biomass and the number of dispersed cells, while average aggregate volume remained lower than WT (Figure 5A–C).

**Figure 5.**
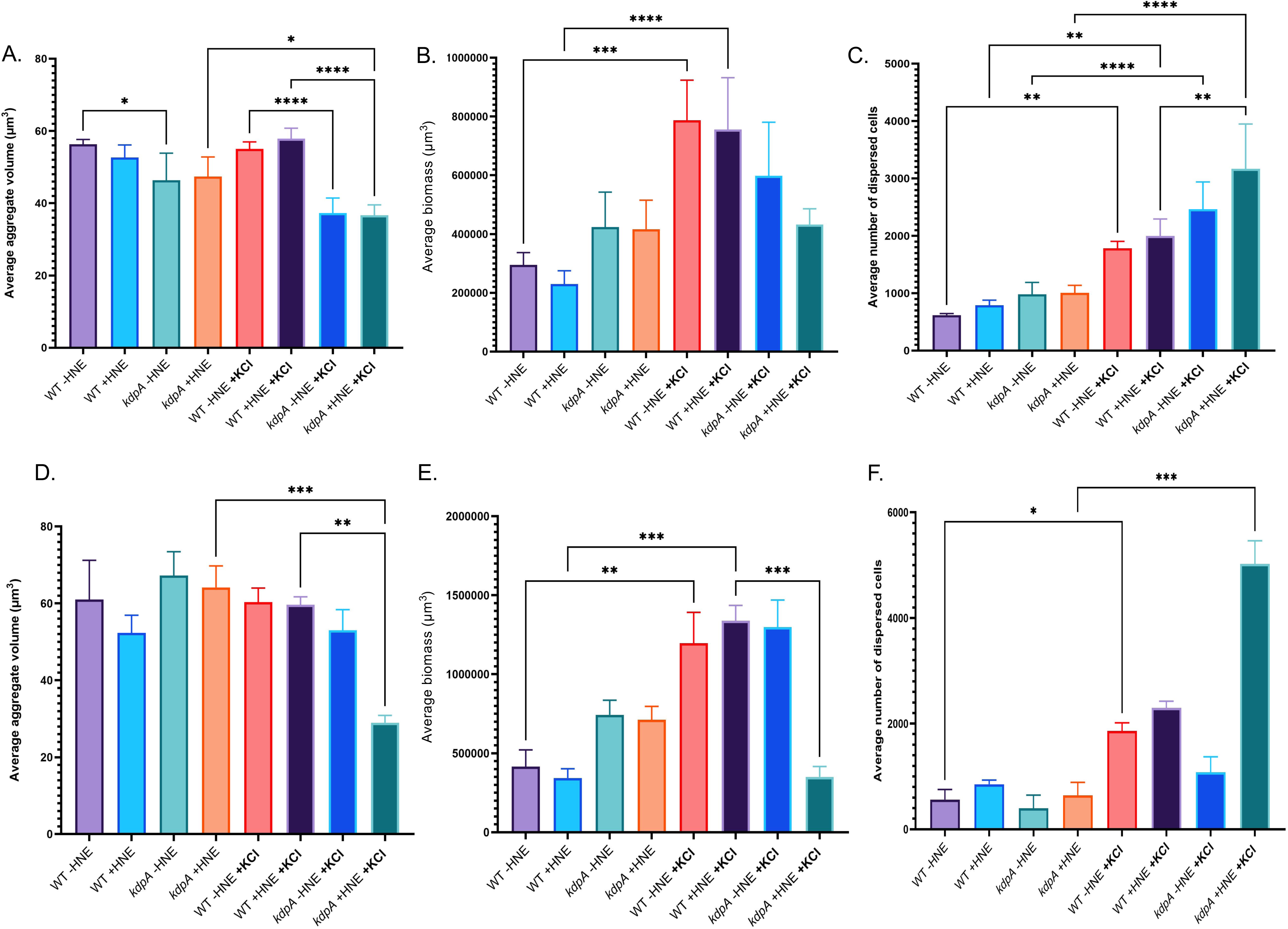
Potassium supplementation modifies kdpA-dependent aggregate phenotypes. WT and kdpA mutant populations were grown in SCFM2 in the absence or presence of 25 mM KCl, with or without 2X HNE (20 μg/L). KCl was added at inoculation, and HNE was added following 4 h of aggregate formation. (A–C) Time-averaged aggregate phenotypes, calculated as the mean of all measurements from each biological replicate across the entire imaging period: (A) average aggregate volume, (B) average biomass, and (C) average number of dispersed cells. (D–F) Aggregate phenotypes at the final time point following HNE exposure: (D) average aggregate volume, (E) average biomass, and (F) average number of dispersed cells. Error bars represent ± SEM. Statistical comparisons for A-C were performed using a two-way ANOVA with main effects only with Sidaks’s multiple comparisons test. Statistical comparisons for D-F were performed using with Krystal-Wallis test. Significance is indicated by an asterisk. Data is representative of three independent biological replicates.

During HNE exposure, KCl supplementation also altered *kdpA* mutant aggregate phenotypes (Figure 5D–F). Aggregate volume was reduced in the KCl-supplemented mutant relative to WT populations (Figure 5D), while total biomass and dispersed-cell number increased (Figure 5E–F). Thus, increased extracellular potassium did not uniformly restore individual mutant phenotypes to WT values. Additionally, KCl supplementation did not fully restore DiBAC signal to WT levels regardless of HNE exposure (Supplementary figure 1A-B).

Analysis of dispersal over time further demonstrated a potassium-dependent change in mutant behavior (Supplemental Figure 5). KCl supplementation altered the temporal dispersal profile of the *kdpA* mutant, with the most pronounced differences evident at later time points following HNE exposure. Together, these findings demonstrate that increased extracellular potassium modifies multiple KdpA-dependent aggregate phenotypes, supporting a functional relationship between potassium availability and KdpA-dependent aggregate physiology.

### Kdp-mediated potassium homeostasis contributes to aggregate physiology during host-derived stress

Together, our findings support a model in which Kdp-mediated potassium homeostasis contributes to *Pa* aggregate physiology and becomes increasingly engaged during host-derived stress (Figure 6). HNE exposure induces the kdpFABC high-affinity potassium transport system, while loss of *kdpA* alters aggregate growth and organization, accelerates dispersal, and changes membrane-associated DiBAC signal. Genetic complementation restores key *kdpA*-dependent phenotypes, whereas manipulation of extracellular potassium modifies the mutant phenotype, demonstrating that these aggregate behaviors are potassium-responsive.

**Figure 6.**
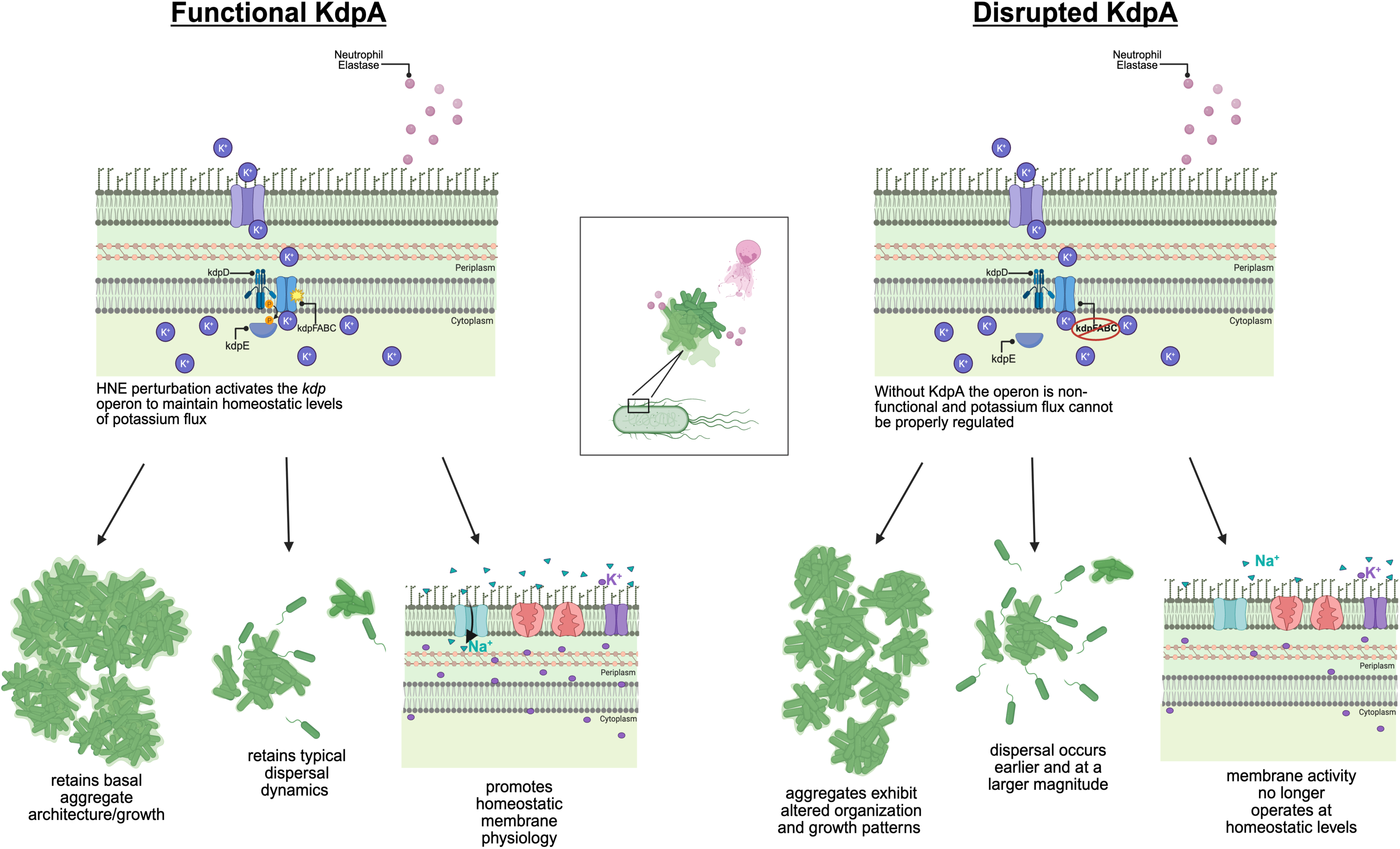
Proposed model for Kdp-mediated potassium homeostasis in *Pa* aggregate physiology during host-derived stress. HNE exposure induces the *kdpFABC* high-affinity potassium transport system, consistent with increased engagement of potassium-homeostasis pathways during host-derived stress. Loss of *kdpA* alters population growth, aggregate organization, dispersal, and membrane-associated DiBAC signal. Genetic complementation restores key *kdpA*-dependent phenotypes, whereas increased extracellular potassium modifies the mutant phenotype, demonstrating potassium responsiveness. We propose that host-derived stress increases engagement of a potassium-homeostasis system that already contributes to aggregate behavior. Intracellular potassium concentrations and potassium flux were not directly measured in this study.

We therefore propose that HNE-associated stress increases engagement of potassium-homeostasis pathways, linking Kdp-dependent transport to the maintenance of aggregate stability. Because intracellular potassium concentrations and potassium flux were not directly measured, ionic perturbation remains a proposed mechanism rather than a demonstrated consequence of HNE exposure.

## DISCUSSION

In this study, we identify potassium homeostasis as a previously underappreciated component of *Pseudomonas aeruginosa* (*Pa*) aggregate physiology. Transcriptomic analysis revealed induction of the high-affinity Kdp potassium transport system following exposure to human neutrophil elastase (HNE), with a greater magnitude of induction in aggregate than planktonic populations (Figure 1C-D). Functional analysis of kdpA subsequently revealed a broader phenotype than an HNE-specific stress response: loss of *kdpA* altered population growth, aggregate organization, dispersal dynamics, and membrane-associated DiBAC signal even in the absence of HNE (Supplementary Figure 2-3). Genetic complementation restored key phenotypes, while co-culture with WT cells did not rescue the mutant despite spatial association between the two populations (Figure 4; Supplementary Figure 4). Manipulation of extracellular potassium further modified multiple *kdpA*-dependent phenotypes (Figure 5; Supplementary Figure 1C-J). Together, these findings suggest that KdpA contributes to the physiological processes that coordinate bacterial growth with multicellular organization and dispersal, while host-derived stress increases engagement of this potassium-homeostasis system.

One of the most striking features of the *kdpA* phenotype was the apparent uncoupling of population growth from aggregate organization. Loss of *kdpA* resulted in increased average biomass while simultaneously reducing average aggregate volume (Figure 2A-B), demonstrating that greater biomass accumulation does not necessarily translate into the formation of larger aggregates. The mutant also exhibited earlier dispersal, suggesting that the effects of *kdpA* extend beyond growth to influence how bacterial populations organize and transition between aggregated and dispersed states. Together, these findings suggest that KdpA contributes to the coordination of population growth, multicellular organization, and dispersal rather than simply regulating bacterial growth itself.

The response to extracellular potassium further supports a functional connection between potassium homeostasis and aggregate behavior. KCl supplementation did not uniformly restore the *kdpA* mutant to a WT phenotype but instead modified aggregate volume, biomass, and dispersal in distinct ways. This argues against a simple model in which increased extracellular potassium merely compensates for loss of the high-affinity Kdp transporter. Rather, the potassium responsiveness of these phenotypes suggests that extracellular potassium availability and Kdp-dependent transport are functionally connected to the regulation of aggregate physiology. Importantly, the differential effects of potassium supplementation on biomass, aggregate volume, and dispersal reinforce the observation that these properties represent related but separable features of aggregate development.

The broader role of KdpA in aggregate physiology provides important context for induction of the Kdp system during HNE exposure. Potassium is a major intracellular cation in bacteria and contributes to fundamental processes including osmotic homeostasis, intracellular pH regulation, and membrane-associated physiology(16, 17). The KdpFABC system provides high-affinity potassium transport and is classically associated with conditions that increase the cellular requirement for potassium uptake(18, 19). Our data do not establish that HNE directly causes intracellular potassium depletion or potassium leakage. However, induction of the Kdp system following HNE exposure suggests that host-derived proteolytic stress increases the requirement for potassium homeostasis. In this model, HNE does not create an entirely new KdpA-dependent phenotype; rather, it increases demand on a homeostatic system that already contributes to normal aggregate growth, organization, and dispersal.

Potassium is increasingly recognized as more than a requirement for cellular homeostasis and can also influence multicellular bacterial behavior. Potassium flux has been shown to mediate electrical signaling within bacterial communities(20), coordinating physiological states across spatially separated populations. In *Pa*, recent work demonstrated that bacteria respond to potassium efflux from airway epithelial cells through the Kdp system, with potassium availability influencing bacterial attachment, coalescence, and biofilm formation(21). Our findings extend this connection between potassium and collective bacterial behavior to free-growing *Pa* aggregates. Rather than affecting a single aggregate property, disruption of *kdpA* altered the relationship between population growth, aggregate organization, and dispersal, while manipulation of extracellular potassium further modified these phenotypes. Together, these observations suggest that potassium availability and potassium transport may contribute to how *Pa* populations organize and transition between multicellular states.

The intersection between potassium homeostasis, host-derived stress, and aggregate behavior may be particularly relevant within the CF airway. Chronic *Pa* infection occurs in a heterogeneous environment characterized by intense neutrophilic inflammation, altered airway chemistry, and substantial spatial variation in nutrient and oxygen availability(23, 24). HNE is abundant in this environment and represents one of several host-derived stresses encountered by bacterial aggregates(10, 25, 26). HNE can also directly influence *Pa* aggregate formation(15), emphasizing that host inflammatory products can shape bacterial multicellular behavior. Rather than suggesting that KdpA functions as an HNE-specific defense mechanism, our findings support a model in which host-derived stress intersects with a core bacterial homeostatic pathway that already contributes to aggregate organization and dispersal. In this context, potassium homeostasis may provide one mechanism through which aggregate populations adapt their organization and dispersal in response to changing host-associated conditions.

Several questions remain regarding the mechanism linking KdpA to aggregate behavior. We did not directly measure intracellular potassium concentrations or potassium flux and therefore cannot determine whether HNE exposure results in potassium depletion or whether the effects of KCl supplementation reflect changes in intracellular potassium homeostasis. Similarly, the altered DiBAC signal observed in the *kdpA* mutant indicates a change in membrane-associated physiology but does not establish a specific change in membrane potential at the single-cell level. Direct measurements of intracellular potassium and membrane potential will therefore be important for defining the physiological basis of the *kdpA* phenotype. It will also be informative to determine whether *kdp* expression varies spatially within aggregates and whether potassium-dependent regulation of organization and dispersal extends to other host-derived or antimicrobial stresses.

Collectively, our findings identify KdpA as a link between potassium homeostasis and *Pa* aggregate behavior. Rather than functioning solely as a response to HNE, KdpA contributes to the coordination of population growth, aggregate organization, and dispersal under basal aggregate-forming conditions, while host-derived stress increases engagement of the Kdp potassium transport system. These findings broaden the role of potassium homeostasis from cellular adaptation to a determinant of multicellular bacterial behavior and provide a framework for understanding how environmental ionic conditions intersect with aggregate physiology during chronic infection.

## MATERIALS AND METHODS

### Strains and growth conditions

*Pa* PAO1 expressing GFP from plasmid pMRP9-1(27) was cultured from frozen stocks overnight in lysogeny broth (LB) at 37°C with shaking at 200 rpm. On the day of each experiment, overnight cultures were diluted 1:5 into fresh LB and grown for approximately 2 h to reach logarithmic phase. Cells were washed with phosphate-buffered saline (PBS; pH 7.0) and inoculated into synthetic cystic fibrosis sputum medium 2 (SCFM2; SyntheBiome, California) at an OD_600_ of 0.05 (∼10^5^ cells/mL). Cultures were incubated statically at 37°C to promote aggregate formation. For co-culture experiments, strains were prepared under the same conditions but were each inoculated at an OD_600_ of 0.025, maintaining a combined starting OD_600_ of 0.05.

### Mutant selection and strain construction

Transcriptomic analysis of *Pa* aggregates exposed to HNE identified PA1633 (*kdpA*) as a candidate gene for further investigation based on its differential expression during HNE exposure. A sequence-verified PAO1 kdpA transposon mutant was obtained from the University of Washington PAO1 ordered transposon mutant library(28). The *kdpA* mutant was transformed by electroporation with the mCherry-expressing plasmid pMP7605(29) to enable visualization by confocal microscopy. The mutant was cultured under the same conditions as the WT strain described above. To confirm that the observed aggregate phenotypes were associated with disruption of *kdpA*, the mutant was genetically complemented with *kdpA*. A pET-21b(+)-based plasmid containing the *kdpA* insert was provided by GenScript and introduced into the *kdpA* mutant by electroporation. The resulting complemented strain was analyzed under the same growth and imaging conditions as the WT and *kdpA* mutant strains.

### Human neutrophil elastase exposure

Lyophilized human neutrophil elastase (HNE; Sigma-Aldrich) was reconstituted according to the manufacturer’s instructions to a stock concentration of 50 μg/L. *Pa* cultures were grown statically in SCFM2 at 37°C for 4 h to allow aggregate formation before HNE exposure. HNE was then added to final concentrations of 5, 10, or 20 μg/L, designated 0.5X, 1X, and 2X HNE, respectively. Unless otherwise indicated, experiments examining the effects of HNE on aggregate phenotypes were performed using 2X HNE (20 μg/L).

### Confocal imaging and image analysis

WT PAO1, the *kdpA* mutant, and the genetically complemented *kdpA* mutant were grown in SCFM2 in the presence or absence of 2X HNE (20 μg/L) and imaged using a Zeiss LSM 880 confocal laser scanning microscope with ZEN image acquisition software. Unless otherwise indicated, experiments were performed with three independent biological replicates, with two technical imaging replicates acquired per biological replicate. Time-lapse imaging was performed for 14 h, with z-stacks acquired at 30-min intervals. Confocal image datasets were analyzed using Imaris 11.0.1 to quantify average biomass, average aggregate volume, and the number of dispersed cells, as previously described(30–34). Quantitative data were subsequently analyzed using RStudio and GraphPad Prism 11.

### RNA preparation and extraction

Non-fluorescent WT *Pa* PAO1 was grown statically in SCFM2 for 4 h at 37°C to allow aggregate formation. HNE was then added to a final concentration of 20 μg/L (2X HNE), and cultures were incubated statically for an additional 6 h. Cultures were subsequently centrifuged, and cell pellets were resuspended in DNA/RNA Shield (Zymo Research), flash-frozen, and stored at −80°C until RNA extraction. For RNA extraction, frozen samples were thawed on ice and transferred to bead-beating tubes (3mm high impact zirconium).

DNase/RNase-free water (100 μL) and lysis buffer (600 μL; Zymo Research) were added to each sample, followed by homogenization at room temperature for 6 min. RNA was purified using the Qiagen RNeasy Mini Kit according to the manufacturer’s instructions and eluted in DNase/RNase-free water. Purified RNA was stored at −80°C. RNA concentration and quality were assessed using the Qubit RNA High Sensitivity and RNA IQ assays (Thermo Fisher Scientific).

### RNA sequencing and differential expression analysis

Purified RNA samples were submitted to Novogene for paired-end 150-bp (PE150) Illumina sequencing. Sequence data were analyzed using the Department of Energy Systems Biology Knowledgebase (KBase)(35). Non-interleaved FASTQ files and the *Pa* PAO1 reference genome were imported into KBase, and biological replicates were grouped by experimental condition using the RNA-seq SampleSet workflow. Raw sequencing reads were assessed for quality using FastQC and aligned to the *P. aeruginosa* PAO1 reference genome using HISAT2. Transcript abundance was quantified using StringTie, and differential gene expression between experimental conditions was determined using DESeq2. Genes with an absolute log2 fold change ≥1.5 and an adjusted significance threshold of α = 0.05 were retained for downstream analysis. Differentially expressed genes were initially screened using a ≥2-fold change threshold. Genes exhibiting <4-fold differential expression were subsequently excluded from further prioritization. To identify transcriptional responses associated with aggregate growth, expression profiles from aggregate populations grown in SCFM2 were compared with those from planktonic populations grown in SCFM. Genes exhibiting strong aggregate-associated differential expression were prioritized for experimental investigation based on the magnitude of the transcriptional response and the availability of corresponding mutants in the PAO1 ordered transposon mutant library. This analysis identified PA1633 (*kdpA*) as a candidate for further investigation.

### DiBAC staining and colocalization analysis

Bis-(1,3-dibutylbarbituric acid) oxonol dyes were obtained from AAT Bioquest as DiBAC_4_(3) (excitation/emission, 493/517 nm) and DiBAC_4_(5) (excitation/emission, 591/615 nm). The two spectrally distinct probes were selected to permit imaging in combination with mCherry-and GFP-expressing *Pa* strains, respectively. DiBAC probes were reconstituted in anhydrous DMSO according to the manufacturer’s instructions and prepared as 1 mg/mL stock aliquots. Following 4 h of static growth in SCFM2, DiBAC was added to bacterial cultures at a final concentration of 5 μg/mL, immediately following HNE addition where indicated. Cultures were incubated in the dark for 30 min before confocal imaging using the Zeiss LSM 880 system described above. Images were acquired at 3-min intervals for 5–6 h and analyzed using Imaris 11.0.1. For colocalization analysis, surfaces were generated independently for detectable DiBAC signal and *Pa* biomass in Imaris. DiBAC surfaces were filtered according to their distance from bacterial surfaces, and DiBAC surfaces with a distance of zero were selected to identify signal spatially associated with bacterial biomass. The proportion of detectable DiBAC signal overlapping with *Pa* biomass was subsequently calculated at 2 and 6 h following HNE addition, corresponding to 6 and 10 h of total aggregate growth, respectively.

### Potassium supplementation

To determine the effect of increased extracellular potassium on aggregate phenotypes, KCl was added to SCFM2 at a final concentration of 25 mM at the time of bacterial inoculation [adapted from (21)]. Cultures were grown statically at 37°C for 4 h to allow aggregate formation, after which 2X HNE (20 μg/L) was added where indicated. Time-lapse confocal imaging was performed using the Zeiss LSM 880 system described above, with images acquired at 3-min intervals for 5–6 h. Image datasets were analyzed using Imaris 11.0.1, and quantitative data were analyzed using RStudio and GraphPad Prism 11.

### Statistical analysis

Statistical analyses were performed using GraphPad Prism 11. Data are presented as mean +/- standard error of the mean (SEM), unless otherwise indicated. Comparisons involving multiple experimental conditions and/or time points were performed using two-way analysis of variance (ANOVA), followed by Šidák’s or Tukey’s multiple-comparisons test, as appropriate. Where assumptions for parametric analysis were not met, comparisons between multiple groups were performed using the Kruskal-Wallis test. Statistical significance was defined as P ≤ 0.05. The statistical test used and number of independent biological replicates (n) for each experiment are specified in the corresponding figure legend.

## ACKNOWLEDGMENTS

We thank Caroline Miller for guidance and support in analyzing RNA-seq data and discussion. We thank other members of the Darch Lab for the discussion and reading of the manuscript.

## AUTHOR CONTRIBUTIONS

Conceptualization and experimental design: S.E.D. and O.M.W. Experiments: O.M.W. Data analysis: S.E.D. and O.M.W. Writing: S.E.D. and O.M.W.

## DATA AVAILABILITY

The RNA-sequencing data generated in this study will be deposited in the NCBI Sequence Read Archive (SRA).

---

**Supplemental Figure 1. Potassium supplementation alters membrane-associated DiBAC signal in *kdpA* mutant aggregates.** (A) Percent overlap between detectable DiBAC signal and aggregate biomass for WT and kdpA mutant populations grown in SCFM2 with or without 25 mM KCl, in the presence or absence of 2X HNE (20 μg/L). Colocalization was quantified in Imaris by determining the proportion of detectable DiBAC signal spatially associated with *P. aeruginosa* biomass. This measurement represents the spatial association of DiBAC signal with bacterial cells and does not represent the percentage of DiBAC-positive cells within the population. (B) Comparison of DiBAC signal overlap with aggregate biomass at 2 and 6 h following HNE addition, corresponding to 6 and 10 h total growth, respectively. Data represent three independent biological replicates. Statistical significance was determined by two-way ANOVA followed by Šidák’s multiple-comparison test; significance is indicated by an asterisk. Error bars represent ± SEM. (C–J) Representative CLSM images showing DiBAC signal associated with WT and kdpA mutant aggregates grown with or without 25 mM KCl and in the presence or absence of 2X HNE (20 μg/L). Scale bars, 10 μm.

**Supplemental Figure 2. Genetic complementation of *kdpA* restores key aggregate growth and dispersal phenotypes.** (A–C) Average aggregate volume (A), average biomass (B), and average number of dispersed cells (C) for WT, kdpA mutant, and genetically complemented *kdpA* mutant populations (indicated by Comp.) grown in the absence or presence of 2X HNE (20 μg/L). Data represent three independent biological replicates. Statistical significance was determined by two-way ANOVA with main effects followed by Šidák’s multiple-comparison test. (D–F) Aggregate volume (D), average biomass (E), and average number of dispersed cells (F) over time for WT, *kdpA* mutant, and complemented *kdpA* mutant populations following addition of HNE. Data represent three biological replicates. Statistical significance was determined by two-way ANOVA followed by Tukey’s multiple-comparison test. Error bars represent ± SEM. Statistical significance is indicated by an asterisk. (G–J) Representative CLSM images of genetically complemented *kdpA* mutant aggregates grown in the absence (G, I) or presence (H, J) of 2X HNE (20 μg/L) at 2 h (G, H) and 6 h (I, J) following HNE addition, corresponding to 6 and 10 h total growth, respectively. Scale bars, 10 μm.

**Supplemental Figure 3. Genetic complementation restores *kdpA*-dependent aggregate phenotypes under non-stressed conditions.** (A–B) Average aggregate volume (A) and average biomass (B) of WT, *kdpA* mutant, and genetically complemented *kdpA* mutant populations (indicated by Comp.) grown in SCFM2 in the absence of HNE. Data represent three biological replicates. Statistical significance was determined by two-way ANOVA with main effects followed by Šidák’s multiple-comparison test. (C–D) Aggregate volume (C) average total biomass (D) at 6 h following the experimental HNE-addition time point (10 h total growth) for WT, *kdpA* mutant, and complemented *kdpA* mutant populations maintained in the absence of HNE. Data represent three biological replicates. Statistical significance was determined using a Kruskal–Wallis test. (E–F) Aggregate volume (E) and average biomass (F) over time for WT, *kdpA* mutant, and complemented *kdpA* mutant populations maintained in the absence of HNE. Data represent three independent biological replicates. Statistical significance was determined by two-way ANOVA followed by Tukey’s multiple-comparison test. Error bars represent ± SEM. Statistical significance is indicated by an asterisk.

**Supplemental Figure 4. Spatial association of wild-type and *kdpA* mutant cells during co-culture.** Percentage overlap between WT and *kdpA* mutant populations grown together in co-culture at 2 and 6 h following addition of 2X HNE (20 μg/L), corresponding to 6 and 10 h total growth, respectively. Percentage overlap represents the spatial association of WT and *kdpA* mutant biomass within co-cultured aggregates. Analysis of individual strain when in co-culture is indicated in bold. Data represent two independent biological replicates. Statistical significance is indicated by an asterisk and was determined by two-way ANOVA followed by Šidák’s multiple-comparison test. Error bars represent ± SEM.

**Supplemental Figure 5. Potassium supplementation alters dispersal dynamics of the kdpA mutant.** WT and kdpA mutant populations were grown in SCFM2 in the absence or presence of 25 mM KCl. KCl was added at inoculation and 2X HNE (20 μg/L) was added following 4 h of aggregate formation. The number of dispersed cells was quantified over the subsequent imaging period to assess the temporal effects of potassium supplementation on aggregate dispersal. Data are presented as mean ± SEM. Statistical comparisons between *kdpA* mutant populations grown with and without KCl were performed at the indicated time points using a two-way ANOVA followed by Tukey’s multiple-comparison test with error bars indicating SEM; significance is indicated by an asterisk. Data represent 3 independent biological replicates.

